# The race goes on: A fall armyworm-resistant maize inbred line influences insect oral secretion elicitation activity and nullifies herbivore suppression of plant defense

**DOI:** 10.1101/2021.05.17.444416

**Authors:** Saif ul Malook, Xiao-Feng Liu, Wende Liu, Jinfeng Qi, Shaoqun Zhou

## Abstract

- Fall armyworm (*Spodoptera frugiperda*) is an invasive lepidopteran pest with strong feeding preference towards maize (*Zea mays*). Its success on maize is facilitated by a suite of specialized detoxification and manipulation mechanisms that curtail host plant defense responses.
- In this study, we identified a Chinese maize inbred line Xi502 that was able to mount effective defense in response to fall armyworm attack. Comparative transcriptomics analyses, phytohormonal measurements, and targeted benzoxazinoid quantification consistently demonstrate significant inducible defense responses in Xi502, but not in the susceptible reference inbred line B73.
- In 24 hours, fall armyworm larvae feeding on B73 showed accelerated maturation-oriented transcriptomic responses and more changes in detoxification gene expression compared to their Xi502-fed sibling. Interestingly, oral secretions collected from larvae fed on B73 and Xi502 leaves demonstrated distinct elicitation activity when applied on either host genotypes, suggesting that variation in both insect oral secretion composition and host plant alleles could influence plant defense response.
- These results revealed host plant adaptation towards counter-defense mechanisms in a specialist insect herbivore, adding yet another layer to the evolutionary arms race between maize and fall armyworm. This could facilitate future investigation into the molecular mechanisms in this globally important crop-pest interaction system.

## Introduction

Fall armyworm (*Spodoptera frugiperda*, FAW) is a lepidopteran herbivore with strong feeding preference towards maize (*Zea mays*). Since its spread from the tropical/sub-tropical region of the New World into Africa and Asia, it has become a serious threat to local maize production and food security (Goergen *et al*., 2016; Sun *et al*., 2021). Management of FAW in its native range of North America is mostly achieved by widespread adoption of transgenic maize cultivars expressing a pyramid of *Bacillus thuringiensis (Bt)* toxic proteins, whereas chemical pesticides and well-designed agronomic practices such as the push-pull system has been deployed in areas with restricted access to transgenic crops (Buntin *et al*., 2004; Midega *et al*., 2018; Yang *et al*., 2021). With accumulation of field-evolved resistance against *Bt* toxins among FAW populations, and the expanding range of this invasive pest, additional control measures are required to device an effective and sustainable pest management strategy for FAW (Zhu *et al*., 2019). Development and adoption of genetically pest-resistance crop cultivars is an important component of a successful integrative pest management system. In maize, a crop well-known for its vast within-species natural variation, FAW-resistant inbred line Mp708 has been developed in 1990, and subsequent mechanistic studies have led to the characterization of an insecticidal cysteine proteinase, Mir1-CP (Williams *et al*., 1990; Jiang *et al*., 1995; Pechan *et al*., 1999; Pechan *et al*., 2000; Pechan *et al*., 2002). Since then, a handful of potential FAW-resistant maize cultivars have been identified through field tests over the years, but few has been further examined for its resistance mechanism (Abel *et al*., 2000; Ni *et al*., 2011; Ni *et al*., 2012; Ni *et al*., 2013; Farias *et al*., 2014)

Meanwhile under laboratory and greenhouse conditions, FAW has emerged as an interesting model species for its specialized dietary preference towards maize compared to its sister *Spodoptera* species. Maize and a few other poaceous crops produce benzoxazinoid compounds as their main specialized metabolites (Zhou *et al*., 2018). These compounds are important for maize defense against diverse phytopathogens and herbivorous insect of different feeding guilds (Ahmad *et al*., 2011; Meihls *et al*., 2013). Glauser *et al*. (2011) found that FAW larvae were not deterred by a common maize benzoxazinoid compound 2,4-dihydroxy-7-methoxy-1,4-benzoxazin-3-one (DIMBOA), and their growth were not inhibited by this compound toxic to other generalist *Spodoptera* larvae. Subsequent studies have demonstrated that FAW was capable of re-glycosylating DIMBOA as well as its toxic degradation product through specialized UDP-glycosyltransferases (UGTs), deactivating this plant defensive compound (Maag *et al*., 2014; Wouters *et al*., 2014; Israni *et al*., 2020). This benzoxazinoid detoxification pathway is likely a part of a broader plant defense tolerance mechanism of FAW, as other insect gene families including cytochrome P450-dependent monooxygenases (CYP450), ATP-binding cassette-containing (ABC) transporters, and glutathione-*S*-transferases (GSTs) are also commonly associated with plant defense tolerance across diverse herbivorous insect species (Kennedy & Tierney, 2013). Most recently, two FAW ABC transporters are reported to be involved in detoxification of *Bt* toxins (Jin *et al*., 2021). Yet, besides these isolated examples, little is known about how FAW respond to feeding on maize genotypes of contrasting resistance phenotypes on a systematic level.

In addition to detoxifying the hallmark defensive metabolite of maize, FAW is also known to manipulate maize inducible defense responses. Feeding by FAW larvae could induce accumulation of toxic benzoxazinoid compounds, ribosome-inactivating proteins, and herbivore-induced plant volatiles (Glauser *et al*., 2011; Chuang *et al*., 2014; De Lange *et al*., 2020). Yet, such induction tends to be weaker than those elicited by sister generalist *Spodoptera* species (De Lange *et al*., 2020). While the exact molecular signaling network that mediate this induction event is yet to be elucidated, it is assumed that this network would be consistent with the theoretical paradigm established in other model plant species. To re-capitulate briefly, herbivore-associated molecular patterns and damage-associated molecular patterns such as volicitins and inceptins would be perceived by plant cell surface receptors. This binding will lead to cross-membrane potential change, calcium ion influx, and activation of mitogen-activated protein kinase (MAPK) signaling cascade. These early signaling events will converge on the induction of jasmonic acid and its bioactive isoleucine conjugate, which will in turn induce the transcriptomic activation of downstream transcription factors and executor genes (Erb & Reymond, 2019). In support of the conservation of this model in maize, *ZmMPK6-silenced* plants demonstrated elevated benzoxazinoids content and enhanced insect resistance (Zhang *et al*., 2021). Therefore, the reduced induction of maize defense responses upon FAW feeding could be a result of an insect-produced signaling interference molecule (*i.e*. an effector), similar to the HARP1 protein first identified in cotton bollworm (Chen *et al*., 2019). Proteomics and targeted metabolomics analyses of FAW saliva have revealed a list of potential plant defense-manipulating effectors including glucose oxidases and diverse phytohormones (Acevedo *et al*., 2017; Acevedo *et al*., 2019). Similarly, a maize-produced chitinase has been identified in the frass of FAW larvae, which could also suppress maize defense responses (Ray *et al*., 2016). Interestingly, the defense-suppressing activity of FAW appeared to be maize-specific as it failed to suppress the defense response of cotton plants (De Lange *et al*., 2020). These observations have led us to hypothesize that maize could evolve counteracting mechanism to nullify the host-adapted defense-suppressing activity of FAW.

In this study, we identified a novel FAW-resistant Chinese maize inbred line, Xi502, through no-choice feeding assay under controlled environmental conditions. Comparative transcriptomic analyses, phytohormone measurements, and benzoxazinoids quantification demonstrated that FAW larvae feeding was not able to suppress the inducible defense response in Xi502 as oppose to the successful defense manipulation in the susceptible maize inbred B73. Parallel transcriptomic analyses of the FAW larvae feeding on these two maize inbreds revealed accelerated transcriptomic re-programing towards maturation when feeding on the susceptible B73 and preferential expression of aromatic compound breakdown-related genes in Xi502-fed larvae. Finally, we demonstrated significant deviation in temporal dynamics of B73 leaf transcriptomic and phytohormonal responses towards FAW oral secretions (OS) collected from feeding on either B73 or Xi502, indicating that the host plant genotype can influence the outcome of maize-FAW interactions both by changing the eliciting activity of FAW OS and by allelic variation in the perceptive components of FAW OS.

## Materials and Methods

### Plant and insect materials

Seeds of diverse maize germplasm were originally obtained from Dr. Jianbin Yan at the Huazhong Agricultural University. Fall armyworms were collected in Yunnan, China and maintained on artificial diet under laboratory conditions over generations by Dr. Yutao Xiao at the Agricultural Genomics Institute at Shenzhen. For all experiments, maize seeds were germinated in 1 L pots filled with commercial potting soil (Pindstrup) with approximately 5 to 1 ratio of vermiculite. Long day light conditions (16 hours) and temperature variation of 24 ± 4°C day, 20 ± 4°C night was provided in the growth chamber. Ten days old maize plants, when the second leaf were fully developed and expanded from the whorl, were used for all experiments.

### Herbivore performance bioassay and elicitation experiments

For inbred line screening, 20 seedlings of each genotype were placed together in a metal-wired cage, and two FAW neonates were placed onto each seedling. After seven days of free-range feeding within the cage, larvae were recovered and weighed. Eight maize genotypes were tested in either round of screening, including B73 as a control for batch effect. Within either batch, larvae weight on the other seven genotypes were compared to that on B73 with Dunnett’s tests. For RNAseq, phytohormone measurement, and benzoxazinoid quantification experiments, one second instar larva was caged onto the second leaf of each seedling with a perforated 45 x 30 x 30 cm^3^ transparent PVC box for designated length of time. All larvae were starved for 2 hours prior to the start of the experiments to ensure prompt start of feeding. Seven to ten independent biological replicates of each genotype and each treatment time points were prepared to account for potential larvae escape or total tissue consumption. Upon harvest, 5 of the biological replicates were weighed, collected into 2 mL centrifuge tubes, and snap frozen in liquid nitrogen for subsequent analyses. For the OS-treated leaf transcriptomics experiment, 3 biological replicates were collected for each treatment and each time point following the same procedure.

### Phytohormone profiling

About 150-200 mg of frozen leaf samples were ground in liquid nitrogen and the powder was extracted with ice-cold ethyl acetate spiked with D6-JA, 13C6-JA-Ile, D5-IAA, 2H6ABA, and 4HSA analytical standards. After centrifugation at 13,000 *g* for 10 min at room temperature (25 °C), supernatants were removed and transferred to fresh centrifuge tubes. The pellets were re-extracted with 0.5 mL of ethyl acetate and centrifuged. Supernatants were combined and then evaporated to dryness on a vacuum concentrator. The residues were resuspended in 0.5 mL of 70% methanol (v/v) and analyzed by HPLC MS/MS (LCMS-8040 system, Shimadzu) according to Wu *et al*. (2007). Target phytohormone concentrations were estimated by ratio of target peaks to their respective analytical standards and normalized by tissue fresh weight.

### Benzoxazinoid extraction

Approximately 150 ± 2.5 mg of frozen grounded leaf samples were used to extract benzoxazinoids. Leaf samples were suspended with MeOH/H2O (50:50, v/v; 0.5% formic acid) in 2 mL centrifuge tubes and vortexed vigorously for 15-20 min. Samples were then centrifuged at 13,000 *g* for 15 min, and 400 μL of the supernatants of benzoxazinoids were transferred to glass vials for analysis on an HPLC MS/MS system (LCMS-8040, Shimadzu) according to Qi *et al*. (2016). Characteristic benzoxazinoid compounds were confirmed with purified standards.

### RNAseq library preparation and data analyses

All RNA samples were extracted following the routine TRIZoL protocol. RNA quality was assessed on 1% agarose gels and with the RNA Nano 6000 Assay Kit of the Bioanalyzer 2100 system (Agilent Technologies, CA, USA). For each sample, 1 μg RNA was used to generate a pair-end sequencing library using NEBNext^®^ UltraTM RNA Library Prep Kit for Illumina^®^ (NEB, USA) following manufacturer’s recommendations and index codes were added to track sequences to each sample, and library quality was re-assessed on the Bioanalyzer 2100 system.

After removing adaptor and low-quality reads, the remaining clean pair-end reads were mapped onto *Zea mays* B73 Refgen v4 or *Spodoptera frugiperda* reference genome with STAR aligner (Dobin *et al*., 2013; Jiao *et al*., 2017; Zhang *et al*., 2020). Raw read counts were calculated with HTseq-count, and differentially expressed genes (DEGs) were calculated with the DESeq2 package with a cutoff of FDR < 0.05 and FC > 2 (Anders *et al*., 2015). Gene ontology enrichment analyses with each set of DEGs were carried out on the agriGO v2.0 online platform, and significantly enriched classes were determined by adjusted p < 0.05 (Tian *et al*., 2017). Raw read counts were converted to fragments per kilobase per million mapped (fpkm) values for principal component analyses (PCA) and pairwise correlation tests between samples in each experiment (Fig S2). In the FAW-attacked maize leaf transcriptomics experiment, one FAW-infested B73 sample showed significant deviation from the remaining 4 biological replicates and was removed from later DEG analyses (Fig. S3).

### FAW OS collection and treatment

FAW larvae were reared on artificial diet until 4-5^th^ instar. Before OS collection, caterpillars were grown on Xi502 or B73 plants for 24 hours. Stork bill forceps were used to gently squeeze the caterpillars to provoke regurgitation, and OS were collected on ice with a pipette and immediately centrifuged to obtain supernatant, which was divided into small aliquots before being stored at −80 °C. Ten days old maize seedlings were wounded by a tracing wheel, and OS or water was applied on the wounding marks and gently rubbed. Treated leaf tissues were harvested as described above at designated time points.

## Results

### Xi502 plants demonstrates stronger resistance against fall armyworms and shorter growth stature

Through caged larvae feeding assay under controlled environmental conditions, we have identified a number of maize genotypes that demonstrated stronger resistance against FAW than the reference maize inbred B73 (Fig. S1). Among these candidates, we further confirmed the resistant phenotype of Xi502 in two separate rounds of experiments, as measured by larvae fresh weight after 7 days feeding (Fig. 1a-c). During these bioassays and later seed propagation work in the field, we noticed that Xi502 plants grew significantly shorter than B73 at both seedling and mature stages (Fig. 1d-f).

**Fig. 1.**
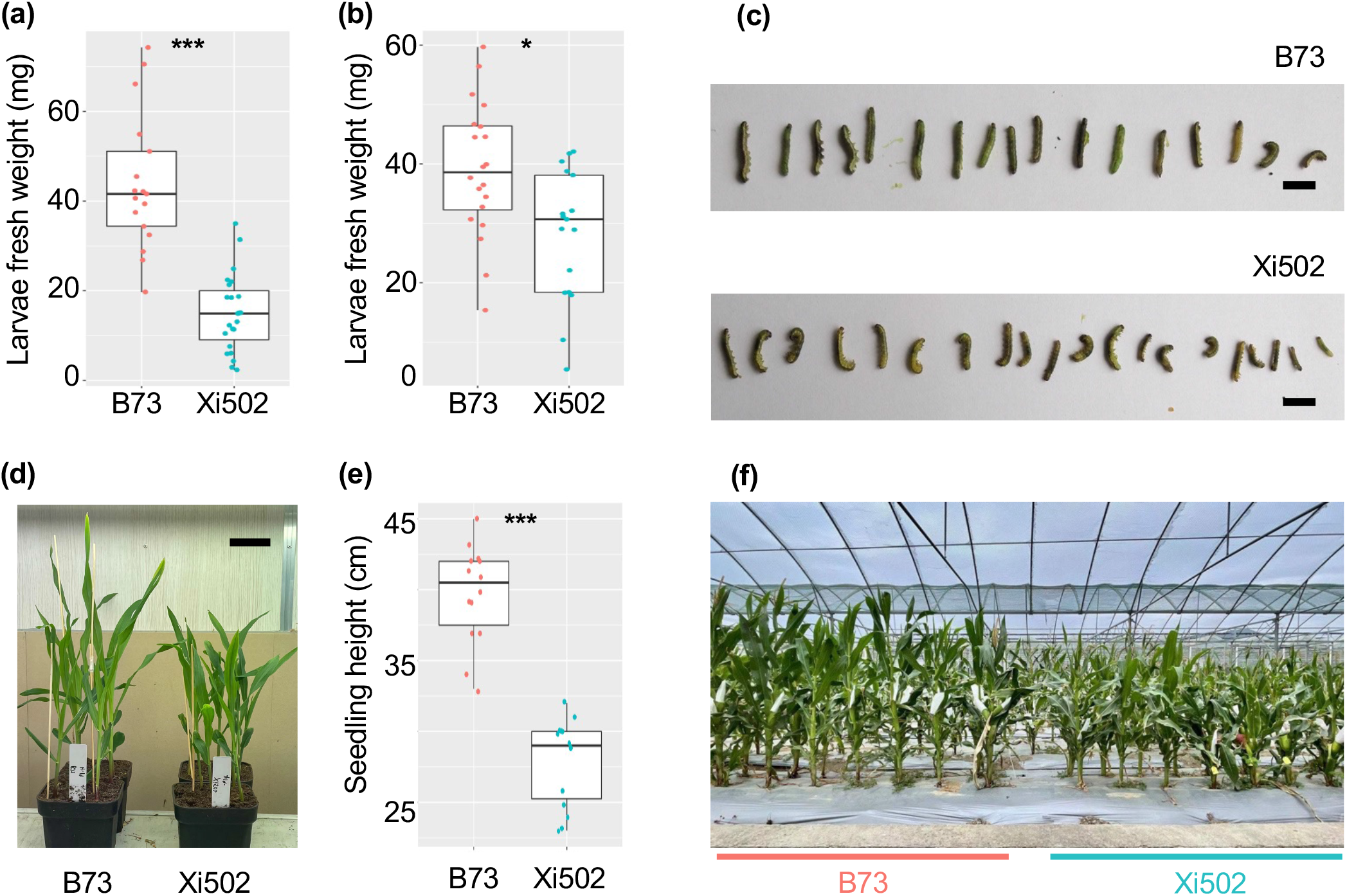
Phenotypic differences between B73 and Xi502. (a, b) In two independent bioassays, FAW larvae grow significantly smaller on Xi502 than on B73 (*p < 0.05; ***p < 0.005; Student’s *t*-tests). The range, quartiles, and mean of either group are shown by the box-and-whisker plots, and the measurement of each larva is represented by a jitter dot. (c) Photographs of snap frozen FAW larvae corpses from the first round of bioassay (Scale bar = 1 cm). (d, e) B73 seedlings grow significantly taller than Xi502 at two-weeks post-sowing (***p < 0.005; Student’s *t*-test; Scale bar = 2 cm). (f) Mature B73 plants grow taller than Xi502 at anthesis stage.

### Fall armyworm feeding induces more pronounced defensive transcriptomic signatures in Xi502 than in B73

Since the FAW resistance mechanism has rarely been studied in Chinese maize genotypes, we further compared the responses of Xi502 and B73 to FAW infestation through whole transcriptome profiling. To that end, B73 and Xi502 leaf tissues subjected to first instar larvae feeding for 24 hours, alongside with empty cage control samples, were collected for RNAseq. Principal component analysis (PCA) of the resulting expression matrix revealed greater separation between FAW-attacked Xi502 and control Xi502 samples compared to their corresponding B73 groups (Fig. 2a; Table S1). Consistently, more than 2,000 DEGs (up-regulated: 1,555; down-regulated: 525) were identified between FAW-attacked and control Xi502, whereas only 175 DEGs (up-regulated: 124; down-regulated: 51) were found in B73 tissues after 24 hours of FAW feeding (Fig. 2b). Among these DEGs, 80 genes were differentially expressed in the same direction in both B73 and Xi502, 95 genes were uniquely induced or suppressed in B73, and 2,000 genes were only affected by FAW feeding in Xi502 (Fig. 2c).

**Fig. 2.**
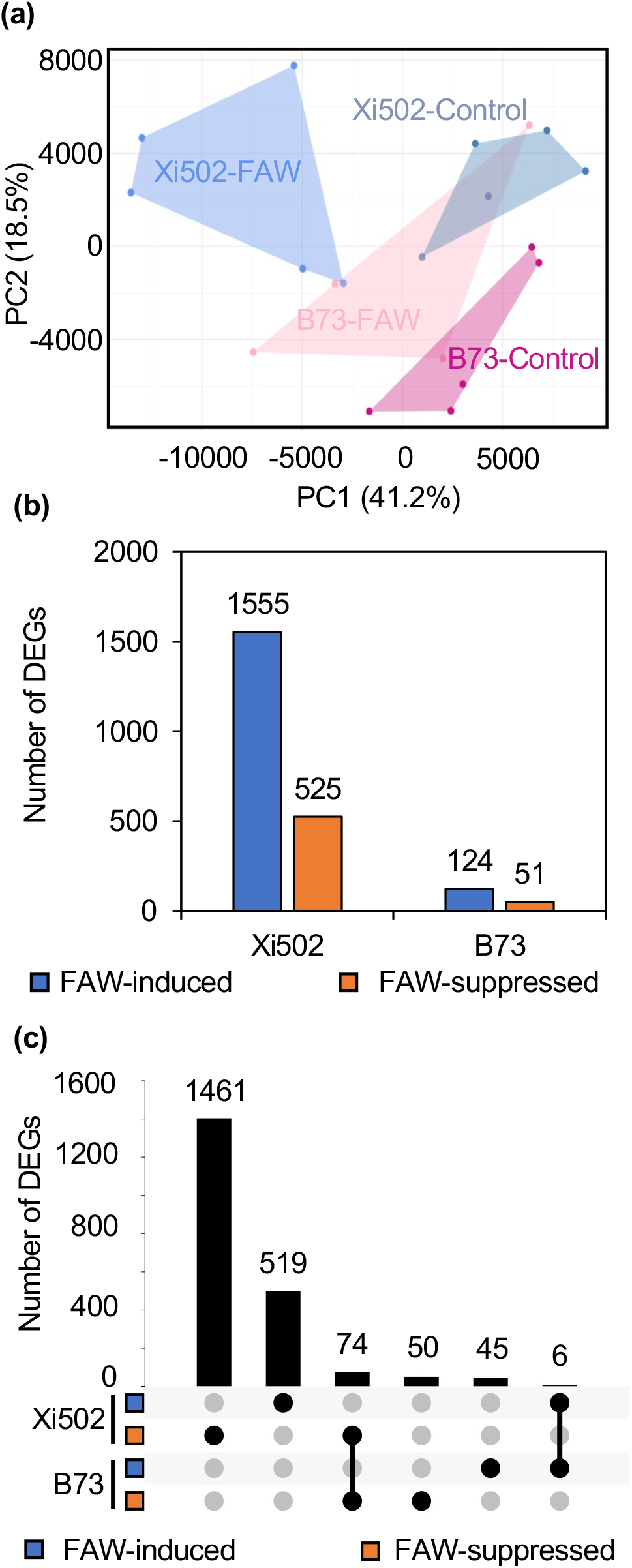
Untargeted comparative transcriptomic analyses of B73 and Xi502 leaves after 24 hours of fall armyworm infestation. (a) PCA result of the RNAseq data from control and FAW-attacked B73 and Xi502 leaf tissue. (b) Summary of DEGs in Xi502 and B73 upon FAW attack. (c) Summary of shared and group-specific DEGs.

Given the small number of DEGs in B73, subsequent gene ontology (GO) term enrichment analyses were done by genotype and by direction of change in expression without looking for overlap between genotypes. Analyses with FAW-induced DEGs in Xi502 showed significant over-representation of genes involved in wounding response (GO: 0009611), response to jasmonic acid (GO: 0009753), and many other categories typically associated with defense against chewing insect herbivores (Table S2). By contrast, the small number of up-regulated genes in B73 were only slightly enriched towards the phenylpropanoid metabolic process (GO:0009698) and endopeptidase inhibitor activity (GO: 0010951; Table S2). FAW-suppressed genes in Xi502 were primarily enriched in photosynthesis-related GO categories, while none of the same category was over-represented by the 51 FAW-suppressed B73 genes (Table S2). Results from these untargeted analyses suggest that Xi502 can mount a more pronounced plant defense response upon FAW attack than B73, including activation of classical stress-related phytohormone signaling pathways and suppression of photosynthetic activities.

Plant-derived toxic proteins and specialized metabolites are known to involve in maize defense against FAW. Yet, two maize defense proteins previously associated with FAW resistance, Mir1-CP (Zm00001d036542) and RIP2 (Zm00001d010371), showed very low level of expression in both genotypes with or without FAW feeding, suggesting that they may not explain the differential resistance phenotypes observed between B73 and Xi502 (Table S1). Another recently identified maize protein potentially enhancing FAW resistance (Zm00001d048950; (Dowd *et al*., 2020) showed stronger induction (as well as higher expression) in B73 compared to Xi502, which was inconsistent with our phenotypic observations (Table S1). Yet, a couple of generic defense-related proteins that have not been specifically associated with defense against FAW, demonstrated higher expression and/or inducibility in Xi502 than B73 (Fig. 3a; Table S1).

**Fig. 3.**
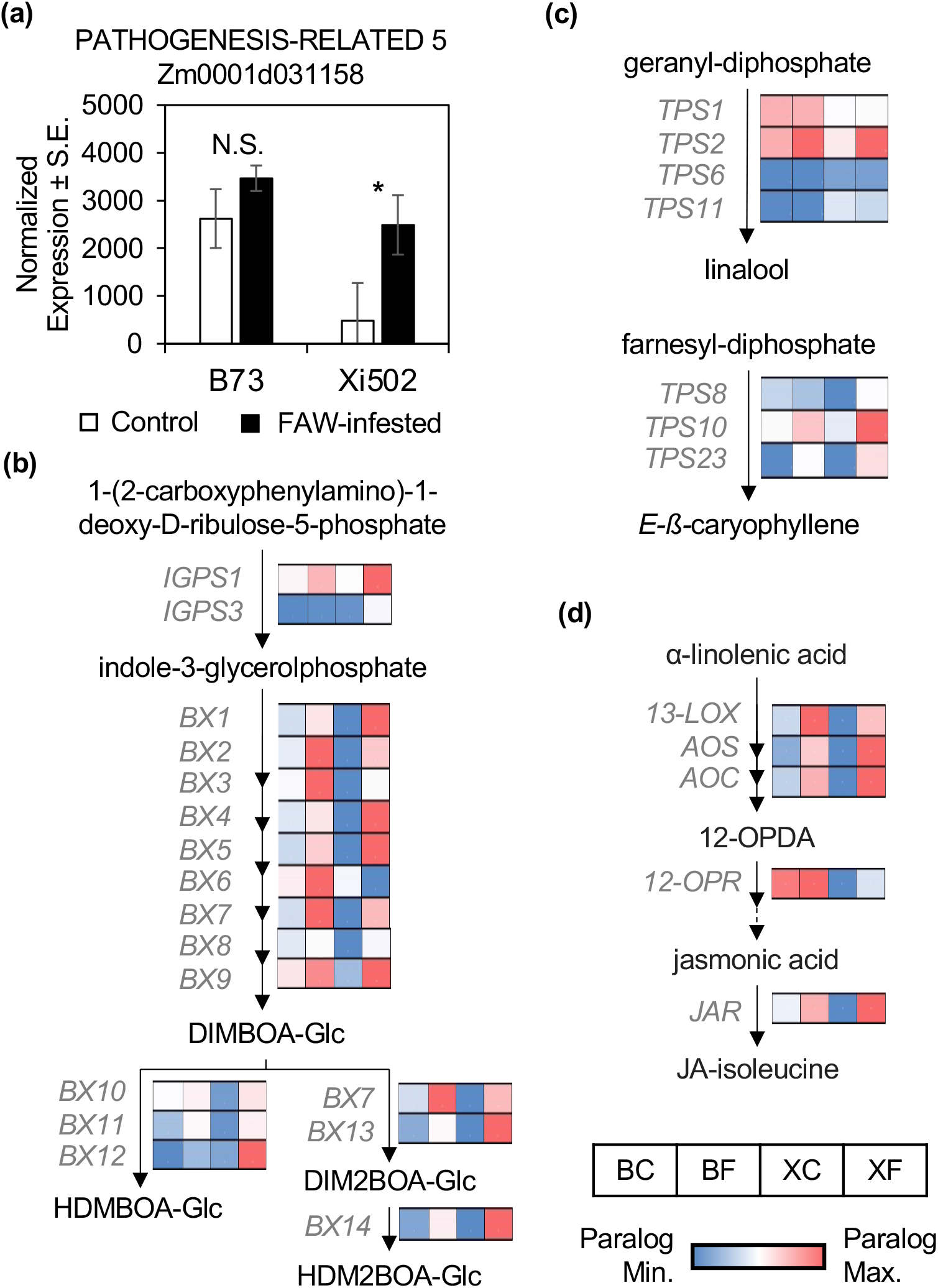
Targeted comparative transcriptomic analyses of B73 and Xi502 upon FAW attack. (a) The maize *PATHOGENESIS-RELATED5* gene is only significantly induced by FAW attack in Xi502 (*FDR < 0.05). N.S. = not significant; S.E. = standard errors. Normalized expression of benzoxazinoid (b), terpenoid (c), and jasmonic acid (d) biosynthetic genes in B73-control (BC), B73-FAW (BF), Xi502-control (XC), and Xi502-FAW (XF) are shown in a blue-to-red color scale. Note the range of the color scale extends from the minimum to the maximum of each set of functionally redundant paralogs (*e.g. IGPS1/3; BX8/9; BX10-12; TPS1/2/6/11; TPS8/10/23*). For jasmonic acid biosynthetic genes, only the paralog with the highest median normalized expression is visualized for simplicity purpose.

In addition to toxic proteins, maize also produces a suite of anti-herbivore compounds, including benzoxazinoids and terpenoids. Constitutively, core benzoxazinoid biosynthetic genes (*i.e. Bx1-9*) express up to more than 10-fold higher in B73 than in Xi502 (as in the case for *Bx2*). Yet, after 24 hours of FAW feeding, the expression of Bx genes reach similar, if not higher, level in Xi502 than in B73, suggesting much stronger inducibility in Xi502. Two indole-3-glycerol phosphate synthases paralogs, IGPS1 and IGPS3, that were recently added to the beginning of the benzoxazinoid biosynthetic pathway also showed comparable constitutive expression level in the two maize genotypes, and significantly higher expression in Xi502 than B73 after FAW feeding (Richter *et al*., 2021). For the more derived steps of the benzoxazinoid biosynthetic pathway, more dramatic difference in inducibility was observed such that *Bx10-14* expression were induced up to 16.8 folds in B73 but induced over 2,000 folds in Xi502 (Fig. 3b). These later steps are known to convert 2,4-dihydroxy-7-methoxy-1,4-benzoxazin-3-one glucoside (DIMBOA-Glc), the primary benzoxazinoid compound in temperate maize inbred lines that has little inhibitory effect on FAW to its more toxic methylated derivative, 2-hydroxy-4,7-dimethoxy-1,4-benzoxazin-3-one glucoside (HDMBOA-Glc). The stronger FAW-inducibility in Xi502 was also observed for terpene synthases responsible for the production of insect parasitoid-attracting linalool (TPS1/2/6/11) and *E*-ß-caryophyllene (TPS8/10/23; Fig. 3c).

Induction of toxic protein and specialized metabolite production are primarily regulated by the jasmonic acid (JA) and ethylene (ET) signaling networks in plants (Erb & Reymond, 2019). Similar to the expression pattern of benzoxazinoid and terpenoid biosynthetic genes, constitutive expression levels of JA metabolic genes were generally higher in B73 than Xi502, but the FAW inducibility tended to be higher in Xi502 (Fig. 3d). None of the ET metabolic gene examined showed significant inducible changes in expression after 24 hours of continuous FAW feeding, though two different paralogs of 1-aminocyclopropane-1-carboxylate oxidases were most highly expressed in B73 and Xi502 (Table S1). Besides small molecule phytohormones, elicitor peptides have emerged more recently as yet another class of important regulators of defense responses in maize. Among the five ZmPROPEP-encoding genes, ZmPROPEP2 (Zm00001d026405) showed highest expression in all of our samples, but no significant difference was found between genotypes or in response to FAW attack (Table S1). On the other hand, ZmPROPEP3 (Zm00001d002138) expression was almost 100-fold higher in Xi502 than B73 under both constitutive and induced conditions (Table S1). Since this peptide has been recently demonstrated to have the strongest elicitation effect, the relatively low absolute expression level (compared to ZmPROPEP2) may be sufficient to induce significant defense responses nevertheless (Poretsky *et al*., 2020).

### Fall armyworm feeding induces stronger and more sustained jasmonates accumulation in Xi502 than B73

Targeted and untargeted transcriptomic analyses consistently suggested that FAW feeding induced stronger defense responses in Xi502 than in B73. Since JA accumulation could occur within a few hours of external stimuli, we conducted a separate FAW feeding time course experiment to measure JA and bioactive JA-isoleucine (JA-Ile) levels in FAW-attacked B73 and Xi502 leaf tissues after 0, 2, 6, or 24 hours of feeding. Using two-way ANOVA with plant genotype and feeding duration as two independent variables, we found only significant elevation of JA and JA-Ile levels in Xi502 tissues after 24 hours of FAW feeding, consistent with the stronger FAW inducibility of JA-biosynthetic genes in this inbred (Table S3). We then compared JA and JA-Ile levels between B73 and Xi502 at each time point with independent Student’s *t*-tests. By this less stringent comparison scheme, we were able to find that Xi502 contained higher level of JA than B73 constitutively as well as after 24 hours of FAW feeding. JA-Ile content was also higher in Xi502 after 24 hours of FAW infestation. After 2 hours of feeding though, Xi502 tissue accumulated significantly higher level of JA, whereas B73 tissue contained higher level of JA-Ile. When comparing measurement at each time point to the mock treatment control within either genotype, JA level was not significantly elevated in B73 until after 6 hours of feeding, while Xi502 had a clear early induction peak after just 2 hours of FAW feeding. JA-Ile level was weakly (but significantly) induced in B73 after 2 hours of feeding, and remained stable at later examined time points. On the other hand, induction of JA-Ile in Xi502 followed a linear increase pattern within the 24 hours testing period, and eventually reached more than 7-fold higher than B73 (Fig. 4a,b).

**Fig. 4.**
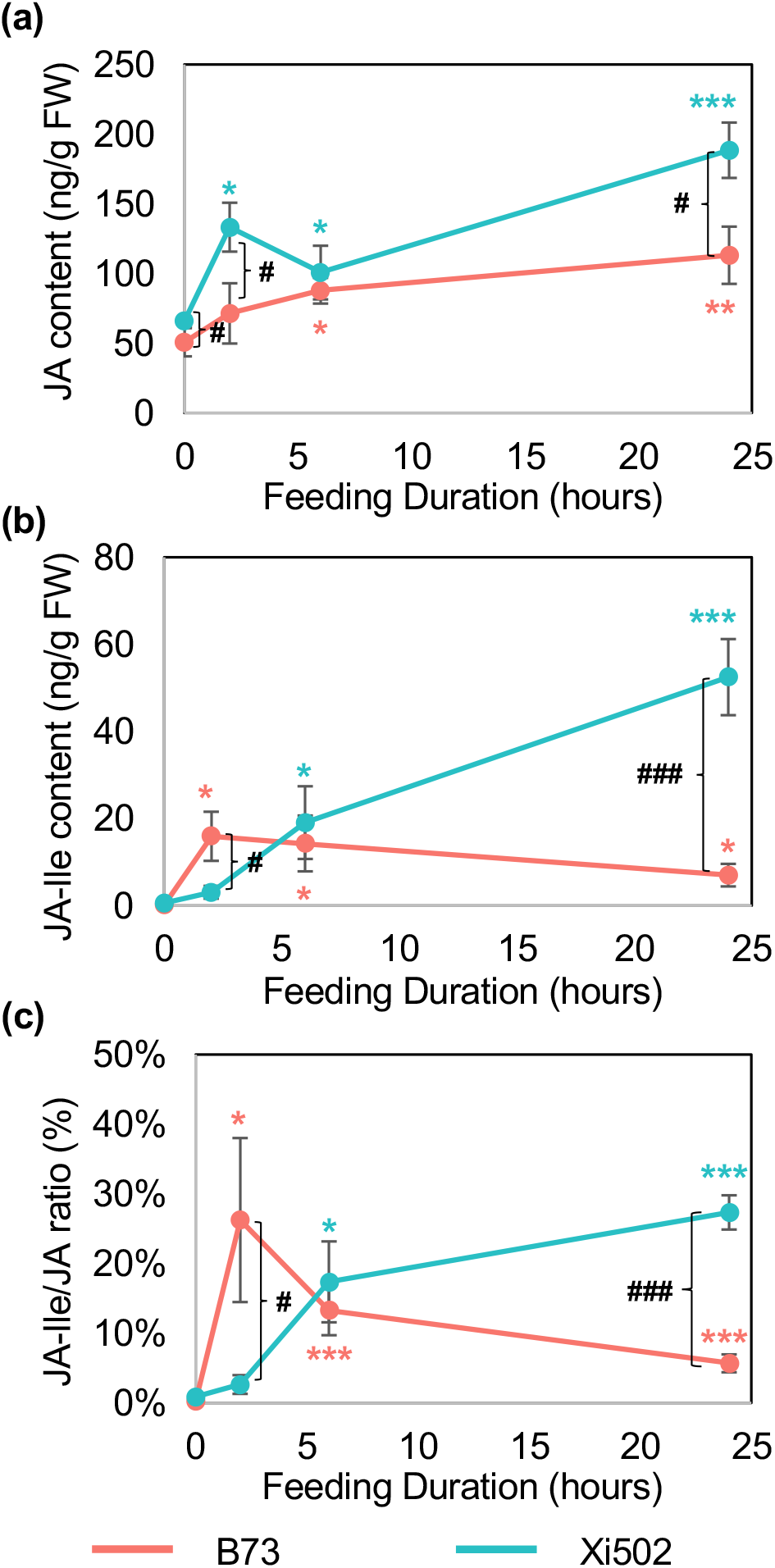
Phytohormone dynamics of B73 and Xi502 upon FAW attack. Measurement of jasmonic acid (JA, a), jasmonate-isoleucine (JA-Ile, b), and their calculated ratio (c) in B73 (orange) and Xi502 (cyan) seedling leaves after 0, 2, 6, or 24 hours of FAW feeding. Significant differences from the 0 hour feeding control of either genotype are indicated by asterisks of their respective color code (*p < 0.05; **p < 0.01; ***p < 0.005; Student’s *t*-tests). Significant differences between the two genotypes at each time point are indicated by pound signs (# p < 0.05; ### p < 0.005; Student’s *t*-tests). N = 5 for each time point and each genotype; error bars = standard errors.

The different post-feeding dynamics of JA and JA-Ile between B73 and Xi502 prompted us to further examine the ratio of these two compounds at each time points. Remarkably, the JA-Ile/JA ratio increased over 270 folds after 2-hours of FAW feeding in B73, but gradually decayed back to about 20% of peak level by 24 hours. By contrast, the JA-Ile/JA ratio demonstrated a slow but constant elevation pattern in Xi502 during FAW feeding, and after 24 hours, reached a comparable level observed in B73 after 2 hours of FAW feeding (Fig. 4c).

### Xi502 accumulates higher levels of benzoxazinoids effective against fall armyworm than B73

Benzoxazinoids are the most abundant defensive metabolites in maize. Various species of benzoxazinoid compounds, however, demonstrate contrasting efficiency in inhibiting the growth of the maize-specialized FAW larvae. The differential expression of benzoxazinoid biosynthetic genes in B73 and Xi502 with or without FAW infestation suggested that the content of benzoxazinoids compounds may vary significantly between these two genotypes (Fig 3b). Measurement of seven different stable benzoxazinoid glucosides produced at different steps of its well-studied biosynthetic pathway after 48 hours of FAW feeding revealed a strong genotypic influence on these compounds both under constitutive and FAW-induced conditions. Surprisingly, despite of the overall lower constitutive expression of benzoxazinoid biosynthetic genes in Xi502, it contained significantly higher levels of four of the seven benzoxazinoid glucosides, under both constitutive and induced conditions (p < 0.05, two-way ANOVA; Figure 5; Table S4). This included the FAW-toxic HDMBOA-Glc, which was almost 7-fold higher in Xi502 than in B73 constitutively. While the lack of variation in DIBOA-Glc and HDM_2_BOA-Glc levels in our experiment were perhaps best explained by their low concentrations, DIMBOA-Glc remained a clear exception to the general trend such that its level was significantly higher in B73 than Xi502 (p < 0.05, two-way ANOVA; Figure 5; Table S4). Significant differences by FAW treatment were only found for DIMBOA-Glc and HMBOA-Glc, but these differences did not exist between control and FAW-attacked groups of the same maize genotype. Interestingly, HM_2_BOA-Glc level was only induced in Xi502, demonstrating a strong genotype-by-treatment effect (p < 0.05, two-way ANOVA; Figure 5; Table S4). Adding together, Xi502 constitutively produced significantly higher level of total benzoxazinoid glucosides, but FAW infestation did not appear to have any impact on their total concentration in either maize genotype.

**Fig. 5.**
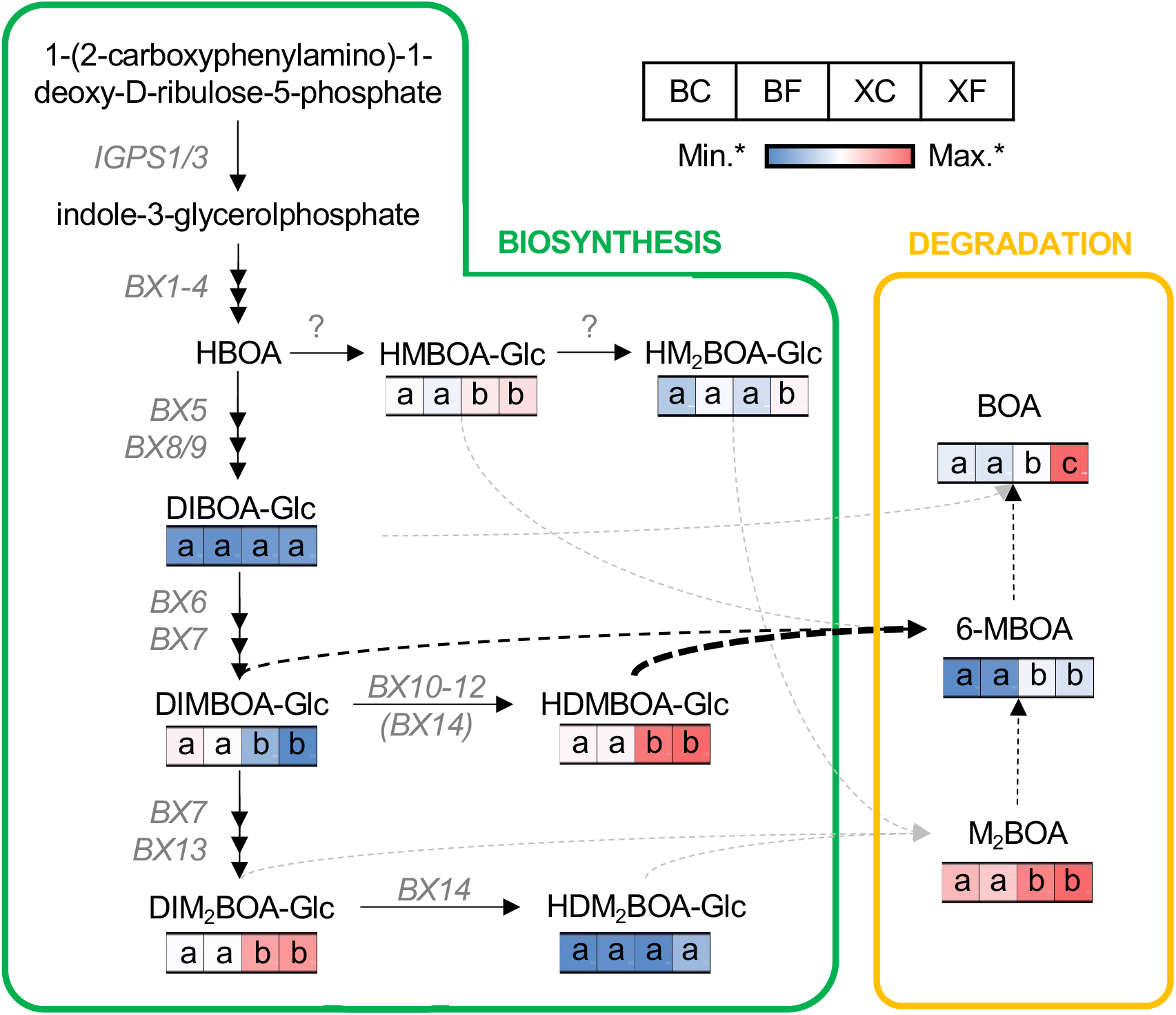
Constitutive and FAW-induced benzoxazinoid content in B73 and Xi502. Concentrations of benzoxazinoids in B73-control (BC), B73-FAW (BF), Xi502-control (XC), and Xi502-FAW (XF) are shown in a blue-to-red color scale. Note that bx compounds produced during biosynthesis (outlined in green) and non-enzymatic degradation (outlined in orange) are plotted on separate scales to better reflect differences between genotype-treatment groups. For each compound and the total benzoxazinoid, significant differences between groups (as determined by two-way ANOVA followed by TukeyHSD) are indicated by different letters in each cell. Known biosynthetic genes are listed at their catalytic steps, with un-confirmed enzymes represented by the question marks. Known and structurally inferred degradation processes are denoted by black and grey dotted arrows, respectively. The faster rate of degradation of HDMBOA-Glc compared to DIMBOA-Glc is reflected by the bolded arrow. BOA: benzoxazolin-2-one; DIBOA-Glc: 2,4-dihydroxy-1,4-benzoxazin-3-one glucoside; DIMBOA-Glc: 2,4-dihydroxy-7-methoxy-1,4-benzoxazin-3-one glucoside; DIM_2_BOA-Glc: 2,4-dihydroxy-7,8-dimethoxy-1,4-benzoxazin-3-one glucoside; HBOA: 2-hydroxy-benzoxazolin-2-one; HDMBOA-Glc: 2-hydroxy-4,7-dimethoxy-1,4-benzoxazin-3-one glucoside; HDM_2_BOA-Glc: 2-dihydroxy-4,7,8-trimethoxy-1,4-benzoxazin-3-one glucoside; HMBOA-Glc: 2-hydroxy-7-methoxy-2H-1,4-benzoxazin-3(4H)-one glucoside; HM_2_BOA-Glc: 2-hydroxy-7,8-dimethoxy-2H-1,4-benzoxazin-3(4H)-one glucoside; 6-MBOA: 6-methoxy-benzoxazolin-2-one; M_2_BOA: 6,7-dimethoxy-benzoxazolin-2-one.

In the same LC-MS assay, we were also able to detect and quantify three benzoxazinoid degradation products, which were believed to actually carry out the anti-feeding bioactivity against lepidopteran herbivores (Zhou *et al*., 2018). Similar to the benzoxazinoid glucosides, all three degradation products were found in significantly higher levels in Xi502. Intriguingly, the final degradation product, BOA, was induced by almost 40 folds only in Xi502 (p < 0.05, two-way ANOVA; Figure 5; Table S4). As a result, the total benzoxazinoid degradation product was constitutively higher in Xi502, and could be further induced in this genotype only.

### Feeding on resistant and susceptible maize tissue leads to distinct transcriptomic response in fall armyworm larvae

As an herbivore with strong preference towards feeding on maize, FAW is known to equip with various counter-defense mechanisms to promote its survival and growth. To examine the potential influence of feeding on the resistant Xi502 and the susceptible B73 on the physiology of FAW, we collected the second instar larvae from the same 24-hours feeding experiment for RNAseq analyses, using artificial diet-fed larvae from the same hatching batch as the control. In sharp contrast to the results from the plant transcriptomic data, PCA of the FAW RNAseq data showed that larvae fed on B73 seedlings had more significant transcriptomic change from the control compared to those feeding on Xi502 (Fig. 6a; Table S5). This result was echoed by the larger number of DEGs in B73-fed FAW larvae (up-regulated: 593; down-regulated: 1,069) than their Xi502-fed siblings (up-regulated: 367; down-regulated: 118; Figure 6b). We found significant overlap between DEGs identified from the two maize diet groups each compared to the artificial diet control, and almost all changes in expression are in the same direction (Fig. 6c). In addition to the shared DEGs, FAW larvae fed on B73 also showed a large number of group-specific DEGs (up-regulated: 751; down-regulated: 498), while DEGs specific to Xi502-fed larvae were considerably fewer (up-regulated: 47; down-regulated: 25).

**Fig. 6.**
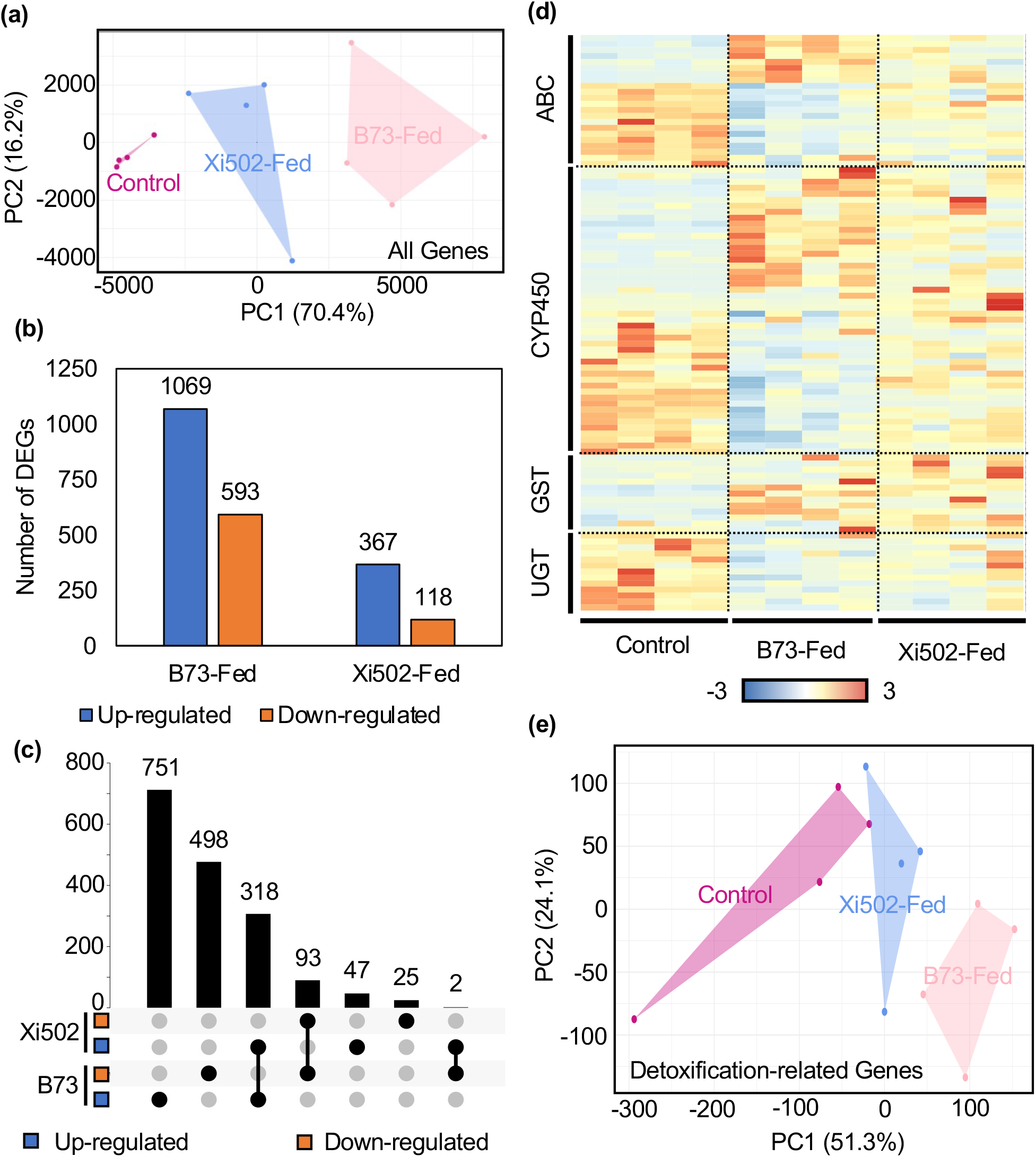
Untargeted comparative transcriptomic analyses of FAW larvae after 24 hours feeding on B73 or Xi502 leaves. (a) PCA result of all expressed genes from FAW larvae feeding on artificial diet (Control), B73 seedlings, or Xi502 seedlings. (b) Summary of DEGs in FAW larvae fed on different diet. (c) Summary of shared and group-specific DEGs. (d) Normalized expression of potential detoxification-related genes in each diet group. ABC: ATP-binding cassette-containing transporters; CYP450: cytochrome P450-dependent mono-oxygenase; GST: glutathione-*S*-transferase; UGT: UDP-glucosyl transferase. (e) PCA results of the potential detoxification-related genes from FAW larvae fed on different diets.

Gene Ontology term enrichment analyses with shared DEGs revealed a number of lipid metabolism-related GO categories were significantly over-represented among up-regulated genes, while cellular transport-related genes were disproportionally down-regulated in the larvae disregard of the maize genotype they fed on (Table S6). Genes up-regulated specifically in B73-fed larvae were functionally-enriched in cytoskeleton organization (GO:0030036), cell adhesion (GO:0007155), and development of reproductive systems (GO:0061458) and sensory perception systems (GO:0007605; Table S6). On the other hand, down-regulated genes specific in B73-fed larvae were over-represented in various metabolic processes including purine nucleotide (GO:0006163), fatty acid biosynthesis (GO:0006633), and alpha-amino acid (GO:1901605; Table S6). Since there were only a few DEGs specific to Xi502-fed FAW larvae, we combined the up- and down-regulated genes for GO term enrichment analysis. Interestingly, genes involved in aromatic compound catabolic process (GO:0019439) was most significantly enriched among these DEGs (Table S6). This GO category was not identified in any of the other four enrichment analyses performed with shared or B73-specific DEGs, suggesting that aromatic compound breakdown may indeed be a specifically-induced physiological process for FAW larvae feeding on Xi502.

To specifically examine the expression dynamics of potential detoxification-related genes, we collected all expressed ABC transporters, CYP450s, GSTs, and UGTs-encoding genes in the FAW transcriptome through Hidden Markov Model search, and compared their expression under the three feeding regimes. Consistent with the PCA results using all expressed genes, PCA of potential detoxification-related genes also demonstrated more significant deviation in B73-fed larvae than in Xi502-fed ones when compared to the artificial diet-fed control group, such that the expression of these genes showed greater change in B73-fed larvae (Fig. 6d,e).

### Oral secretions from fall armyworm feeding on B73 and Xi502 elicit distinct temporal dynamics in B73

Components of FAW OS and frass have been reported to suppress maize defense responses (Acevedo *et al*., 2017; Acevedo *et al*., 2019). Meanwhile, the compositions of FAW OS and frass are known to be significantly influenced by the larvae’s diet, and the very plant defense-suppressing molecules can originate from the consumed plant tissues (Ray *et al*., 2016). The overwhelming difference in B73 and Xi502 leaf tissue response to FAW feeding at transcriptome level led us to hypothesize that the OS resulted from FAW feeding on these two maize genotypes could have distinct biochemical composition and hence differential bioactivity in maize-FAW interaction. To test this hypothesis, OS were collected from FAW larvae reared on B73 or Xi502 leaves (referred to as OS_B73_ and OS_Xi502_ hereafter, respectively), and used to treat B73 leaves after mechanic wounding. Leaf tissues were collected at 15 minutes, 2 hours, and 6 hours post-treatment, with wounding plus water treatment (W+W) as the control for each time point. In support of our hypothesis, principal component analyses of the RNAseq data revealed clear distinction between W+OS_B73_ and W+OS_Xi502_ treated groups at 15 minutes and 2 hours post-treatment, while the differentiation between the OS and water treatment groups was apparent across all three tested time points (Fig. 7a-c; Figure S4; Table S7).

**Fig. 7.**
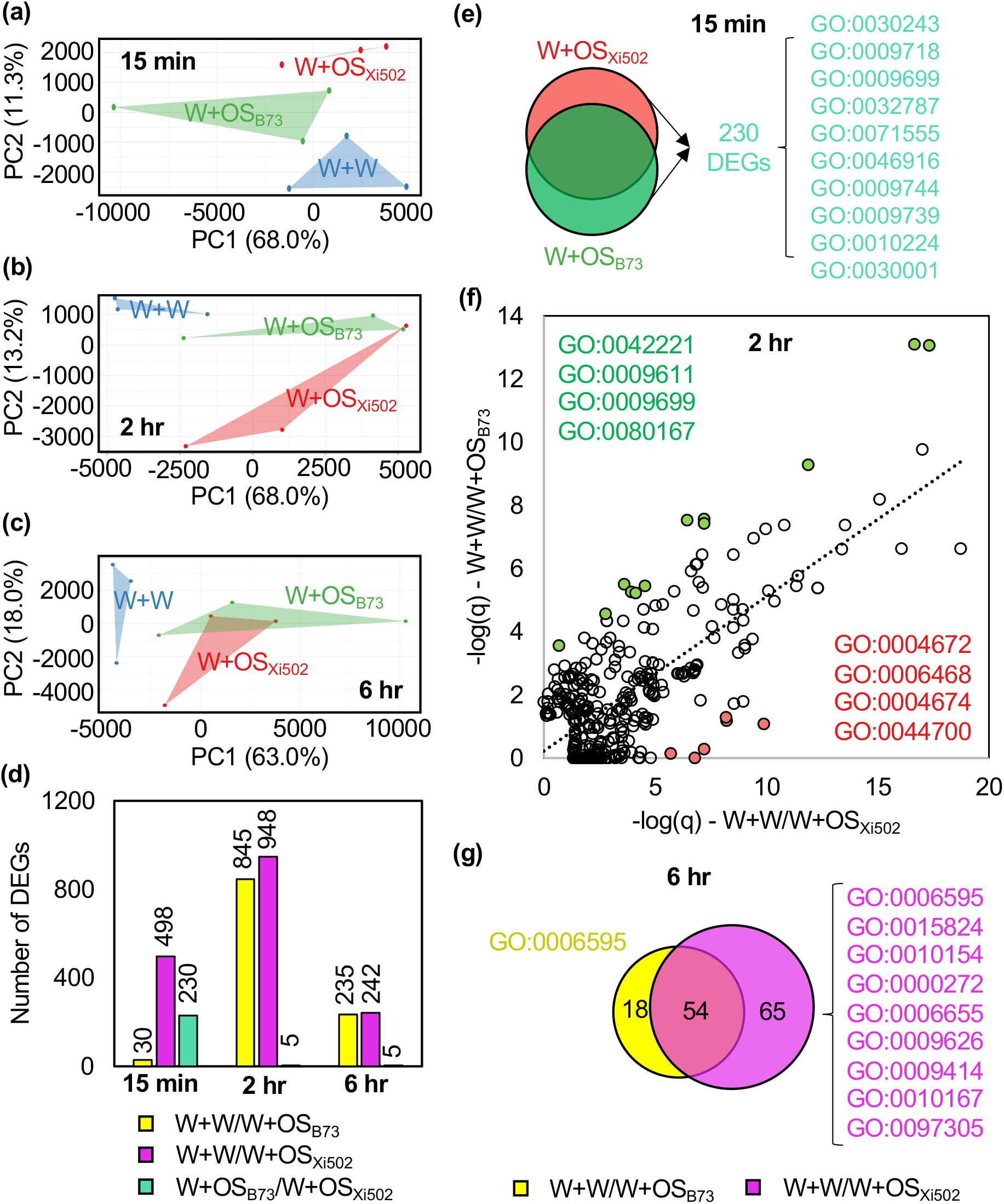
Distinct temporal dynamics in B73 leaf transcriptome elicited by oral secretion collected from FAW fed on B73 and Xi502. (a-c) PCA result of the RNAseq data from B73 leaf tissue treated with wounding and water (W+W), wounding and oral secretion (OS) collected from B73 (W+OS_B73_), or wounding and OS collected from Xi502 (W+OS_Xi502_) collected at 15 minutes (15 min), 2 hours (2 hr), or 6 hours (6 hr) post-treatment. (d) Summary of DEGs in each of the groups above at each time point. (e) Significantly enriched GO terms for DEGs between W+OS_B73_ and W+OS_Xi502_ at 15 minutes posttreatment. (f) Preferentially-enriched GO terms among W+OS_B73_ and W+OS_Xi502_ DEGs at 2 hours posttreatment. (g) Specifically-enriched GO terms among W+OS_B73_ and W+OS_Xi502_ DEGs at 6 hours posttreatment.

In congruence to the PCA results, more DEGs were identified between the W+OS_Xi502_ group and the W+W group than between the W+OS_B73_ group and the W+W group at all three time points (Fig. 7d). When comparing the two OS treatment groups directly, 230 DEGs were found at 15 minutes post-treatment, whereas only 5 genes were differentially expressed between the two groups at either of the two later time points (Fig. 7d). Gene ontology enrichment analyses of the 498 DEGs between the W+OS_Xi502_ group and the W+W group showed that a number stimuli-responsive GO categories, including response to chitin (GO:0010200) and response to jasmonic acid (GO:0009753), were over-represented among these DEGs (Fig. 7d; Table S8). In contrast, no GO term was enriched among the 30 DEGs between the W+OS_B73_ group and the W+W group at the same time point. The 230 DEGs found between the two OS-treated groups were enriched towards a number of categories related to specialized metabolism (*e.g*. anthocyanin-containing compounds, phenylpropanoids), metal ion homeostasis, and responses to various stimuli (*e.g*. sucrose, gibberellin, ultraviolet-B; Figure 7e; Table S8).

Both OS treatments induced a large number of DEGs at the later time points compared to their respective control groups, and GO enrichment analyses with these DEGs resulted in dozens of over-represented categories (Table S8). To identify possible differentially-influenced physiological processes between these two OS treatments, we combined the enriched GO terms from both sets of DEGs identified between W+W/W+OS_B73_ and W+W/W+OS_Xi502_ at either time points, and plotted their -log(q) values from the two comparisons. At 2 hours post-treatment, we observed significant positive correlation (R^2^ = 0.5282) between the two sets of enriched GO terms, suggesting that the comparable number of DEGs from these two comparisons were also mostly involved in similar sets of physiological processes (Fig. 7f). Interestingly, identification of GO terms that significantly deviated from this generally positive trend demonstrated that a number of protein kinase activity and phosphorylation-related terms were preferentially enriched among W+W/W+OS_Xi502_ DEGs (Fig. 7f). At 6 hours post-treatment, we were not able to find the same kind of positive correlation (R^2^ = 0.2749), indicating that these two sets of DEGs, though similar in number, were involved in distinct physiological processes. Indeed, when qualitatively comparing the significantly enriched GO terms from these DEG sets, while almost all GO terms enriched among W+W/W+OS_B73_ DEGs were also found over-represented among W+W/W+OS_Xi502_ DEGs, this latter group of DEGs was also enriched for 65 other GO categories, involved in diverse processes including proline transport, phospholipid metabolism, terpenoid metabolism, and plant-type hypersensitive responses (Fig. 7g).

### Inducible phytohormone dynamics is affected by both the source of oral secretions and the host plant genotype

To further explore the dynamic plant response towards FAW OS collected from different source diet, we treated B73 and Xi502 seedling leaves with water (as control), OS_B73_, or OS_Xi502_ after mechanic wounding. Local treated tissues were collected 2 hours post-treatment, and used for simultaneous quantification of five different phytohormones (Table S9). The epitomic insect defense hormone JA was significantly induced in B73 leaves after OS_Xi502_ but not OS_B73_ treatment. Surprisingly, though the constitutive JA level was significantly higher in Xi502, it was significantly depleted by OS_Xi502_ treatment, while OS_B73_ treatment induced a weaker and insignificant reduction as well (Fig 8a). The similar pattern was also observed for the bioactive JA-Ile conjugate, and the JA-Ile/JA ratio showed no significant difference across the board (Fig 8b,c). Content of another phytohormone that has been extensively associated with defense against chewing herbivores, abscisic acid (ABA), was induced only by OS_B73_ but not OS_Xi502_, though this induction is only statistically significant in B73. By contrast, only OS_Xi502_ treatment could induce significant indole-3-acetic acid accumulation in the two hosts (Fig 8e). Interestingly, salicylic acid (SA), which often plays counteracting role against JA in plant defense responses, was significantly depleted in B73 after OS_Xi502_ treatment and in Xi502 after OS_B73_ treatment, two scenarios that less likely occurring under natural conditions (Fig 8f). In congruence with the results of post-OS-treatment transcriptomics analyses, phytohormone dynamics also clearly support that OS_B73_ and OS_Xi502_ have distinct host response elicitation activity profile. Furthermore, by including both host plant genotypes in this experiment, we were able to demonstrate that B73 and Xi502 could respond differently even towards the same source of OS treatment (as in the cases of JA and JA-Ile), underlying that the host plant response is a function of both the source of OS and the host genotype.

**Fig. 8.**
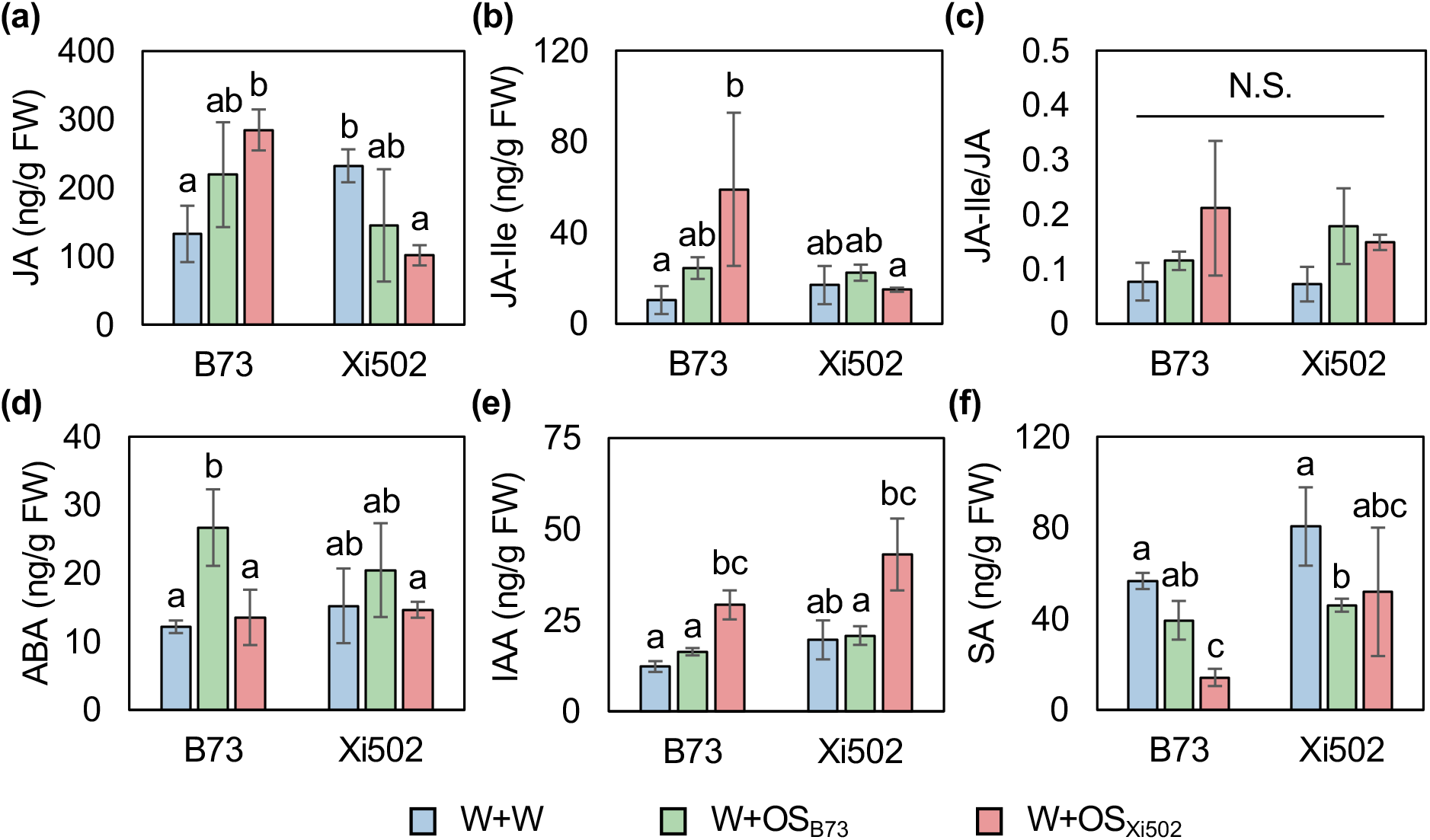
Phytohormone content in B73 and Xi502 leaves under different oral secretion treatment regimes. Concentrations of (a) jasmonic acid (JA), (b) jasmonic acid-isoleucine conjugate (JA-Ile), (c) their calculated ratio, (d) abscisic acid (ABA), (e) indole-3-acetic acid (IAA), and (f) salicylic acid (S_A_) in B73 and Xi502 seedling leaves treated with wounding and water (W+W), wounding and oral secretion (OS) collected from B73 (W+OS_B73_), or wounding and OS collected from Xi502 (W+OS_Xi502_) after two hours. N = 3 for each genotype-treatment group. Groups significantly different from each other according to two-way ANOVA and TukeyHSD (p < 0.05) are denoted by different letters on top of their representative columns. Error bars = standard deviation.

## Discussion

As a successful specialist herbivore, FAW has evolved an arsenal of counter-defense mechanisms including defensive metabolite detoxification and inducible response suppression to promote its own survival and development on the preferred host plant species (Maag *et al*., 2014; Wouters *et al*., 2014; De Lange *et al*., 2020; Israni *et al*., 2020). These specialized counter-defense measures of FAW posed additional challenges to developing genetically FAW-resistant maize cultivars by harnessing the innate maize biochemical defense. Nevertheless, natural variation within maize has led to identification and breeding of maize inbred lines with enhanced FAW resistance, and such resistance has often been attributed to specific defensive biomolecules such as the Mir1-CP protease in Mp708 and the constitutively accumulated FAW-toxic HDMBOA-Glc among tropical maize inbreds (Pechan *et al*., 2000; Meihls *et al*., 2013). However, even in the classic FAW-resistant inbred Mp708, the inducible accumulation of Mir1-CP is not the only trait potentially associated with the heightened resistance phenotype, such that it also has constitutively higher level of jasmonates and volatile terpenoids (Shivaji *et al*., 2010; Smith *et al*., 2012). Indeed, successful defense against insect herbivores require a suite of concerted response as implicated by the transcriptomic and metabolomic dynamic demonstrated here in Xi502, as well as in previous studies on maize attacked by generalist *Spodoptera* species, and one would expect that a singular change on any specific branch of the defense response would either be insufficient to stop the herbivores or impose a strong enough selective pressure that a counteracting mechanism would quickly arise in the insect populations (Fig. 2-5; (Erb *et al*., 2009; Tzin *et al*., 2017). Therefore, the molecular signaling network that regulates the herbivore-inducible responses in plants likely presents an important battleground for the arms race between insect herbivores and their host plants. In support of this hypothesis, we found the largest number of DEGs between the two FAW OS treatment groups at 15 minutes post-treatment when the response has yet to propagate to the downstream executor genes (Fig. 7d,e).

Furthermore, we showed that protein kinase activity and phosphorylation-related GO terms were preferentially enriched in OS_Xi502_ treatment groups at 2 hours post-treatment (Fig. 7f). This observation suggested that OS_Xi502_ could induce a more pronounced activation of the MAPK and/or calcium-dependent protein kinase (CDPK) signaling cascade in maize, which function upstream to the plant JAZ proteins targeted by the HARP1 effector recently identified in cotton bollworm (Chen *et al*., 2019). Since the only variable between these two groups was the genotype of host plants that were used to produce the OS, we speculate that a progenitor plant molecule may be ingested and modified by the FAW larvae, and secreted back into the plant cell as an effector molecule to interfere with the MAPK/CDPK signaling cascade, and the differential elicitation activity between OS_B73_ and OS_Xi502_ could be explained by the presence of modification-resistant progenitor molecule in Xi502. In result, OS_Xi502_ would contain no (or less) functional effector molecules so that the herbivory-inducible signaling cascade could function as norm (Fig. 9a,b). Alternatively, Xi502 may contain a yet unknow inhibitor molecule that suppress the biosynthesis and/or secretion of insect-produced effectors (Fig. 9c,d). Furthermore, the strength of defense signals in this system is not exclusively determined by the eliciting activity of FAW OS, as we demonstrated that different host plant genotypes could behave in contrasting fashions even when treated by the same OS (Fig 8a,b). This would suggest that allelic variation in the hypothetical effector-targeted protein kinases between B73 and Xi502 could also play a role in the differential FAW-inducibility in these two genotypes.

**Fig. 9.**
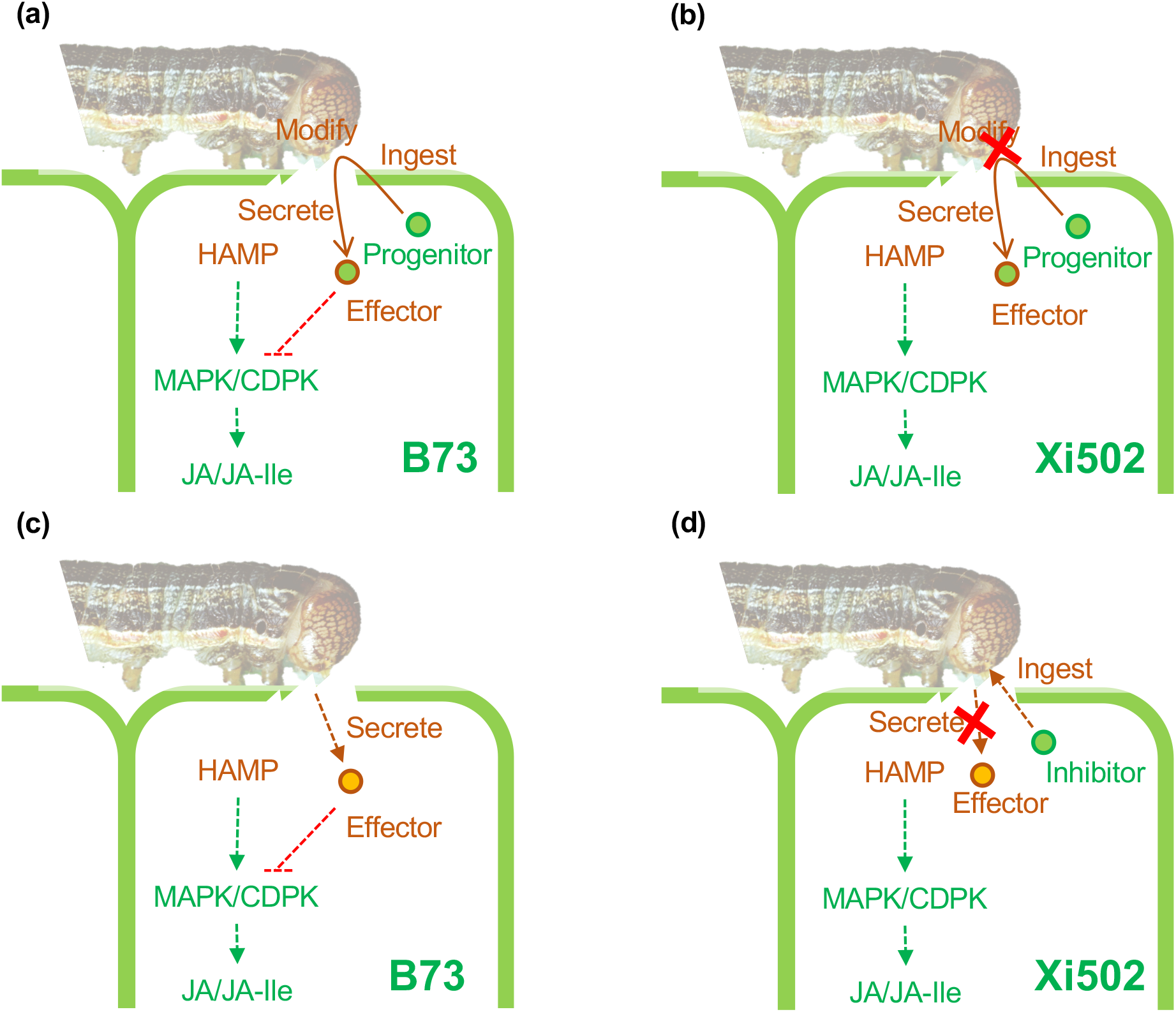
Hypothetical models of host plant genotype’s influence on the elicitation activity of FAW oral secretion. (a) In the progenitor model, a plant progenitor molecule is ingested, modified, and re-secreted into plant cells as an effector to interfere with early activated plant kinases to suppress downstream defense responses in the susceptible B73. In the resistant Xi502, the same plant molecule is resistant to the modification and hence no (or less) effector molecules will be produced, and normal plant defense responses are restored. (b) In the alternative inhibitor model, the effector molecule is directly produced and secreted by the insect, and the resistant Xi502 contains an ingestible inhibitor that hampers the effector biosynthesis and/or secretion processes. In all diagrams, plant- and insect-derived components are shown in green and orange, respectively. HAMP: herbivore-associated molecular pattern; MAPK: mitogen-activated protein kinase; CDPK: calcium-dependent protein kinase; JA: jasmonic acid; JA-Ile: jasmonic acid-isoleucine conjugate.

The initial observation that Xi502 showed elevated resistance against FAW as well as shorter growth stature had prompted consideration of possible growth-defense tradeoff in this particular genotype (Fig. 1). In support of this idea, Xi502 did contain higher constitutive level of total benzoxazinoids and JA than B73 (Fig. 4&5). However, almost all of the defense-related genes we have examined specifically showed higher constitutive expression level in the susceptible B73. The consistent trend was that Xi502 demonstrated stronger defense inducibility upon FAW attack at all of the three aforementioned levels, strongly suggesting that the FAW-resistance phenotype was at least in part due to genetic variation in its herbivore-responsive signaling network (Fig. 3&4). Hence, we can neither support or refute the hypothetical linkage between resistance and growth in Xi502 with these contradicting evidences between gene expression, jasmonates levels, and benzoxazinoids contents. Ideally, this hypothesis could be better tested by examining the FAW-resistance and growth stature phenotypes among the parental, sibling, and segregating filial lines of Xi502.

In addition to multi-level characterization of plant responses to FAW attack, we also examined the transcriptomic responses of the larvae feeding on these two host genotypes with the goal of depicting a more complete picture of this plant-insect interaction system. Though the larvae under different feeding regimes were undiscernible to phenotypic observations after 24 hours of feeding, significant re-programing had readily occurred at the transcriptomic level (Fig. 6). Enrichment towards sensory and reproductive developmental processes among B73-specific up-regulated genes presented clear evidence of better larval development on this susceptible host (Table S6). This interpretation was further supported by the B73-specific down-regulation of various nutrient metabolic processes (Table S6). Unexpectedly, many detoxification-related genes, with the exception of aromatic compound breakdown-related genes, also showed more significant change in expression in B73-fed larvae compared to their Xi502-fed siblings, though the latter group was exposed to much more hostile defensive metabolites in its diet (Fig. 6d,e). This may suggest that Xi502 has evolved mechanisms to suppress the expression of FAW detoxification genes. Host-adapted insect transcriptomic responses have been hypothesized to link with important biological functions such as host manipulation and nutrient assimilation (Petre *et al*., 2020). Among the handful of transcriptomics studies examining herbivore responses on different host plants, the transcriptome dynamics appeared to be greater in chewing herbivores (~5% DEG) than in phloem-suckers (0.1~1% DEG; (Birnbaum *et al*., 2017; Mathers *et al*., 2017; Boulain *et al*., 2019; Tan *et al*., 2019). The 1,662 (7.1%) and 485 (2.1%) DEGs between B73- and Xi502-fed FAW larvae compared to their siblings grown on artificial diet underscored the significant influence of host plant genotype on the herbivore transcriptome plasticity, and in this case, a large number of DEGs were not directly linked to host adaptation but rather involved in developmental processes of the insects themselves, even with the transient feeding period (24 hours) on each host plant (Fig. 6a,b).

Finally, simulation of insect herbivory by wounding and OS treatment is a common practice in plant-insect interaction research (Waterman *et al*., 2019). Our result of differential elicitation activity of FAW OS collected from larvae fed with different host plant genotypes should raise the caution of how the OS used in such simulated herbivory experiments were prepared, and whether this preparation process could introduce any artifact into the results of these experiments. As researchers look further into plant natural variation for sustainable insect pest management solutions, the multi-faceted influences of host genotypic variation on plant-insect interactions must be carefully and thoroughly evaluated.

## Data availability statement

All RNAseq clean reads are uploaded to NCBI online depository in fastq format and accessible under the following BioProjects: PRJNA723461 – FAW-infested maize leaf transcriptomes; PRJNA729598 – FAW larvae transcriptomes; PRJNA730324 – FAW OS-treated maize leaf trancriptomes.

## Acknowledgements

The authors would like to thank Drs. Jianbin Yan and Yutao Xiao for generous supply for plant and insect materials. We would also like to thank Drs. Suhua Li, Xiaowen Shi, Jian Yan, and Georg Jander for constructive suggestions in preparing the manuscript. This work has been financially supported by National Natural Science Foundation of China (Award 32000229 to S.Z. and Award 31772179 to J.Q.), Shenzhen Peacock Plan (KQTD20180411143628272), and Shenzhen Excellent Science & Technology Innovation Talent Project (RCBS20200714114918029).

## Author contributions

This project is conceived by SuM, WL, and SZ. SuM performed bioassays and tissue preparation for omics experiments, and analyzed bioassay results. XFL analyzed transcriptomics data. JQ performed phytohormone and benzoxazinoid measurements and analyzed data from these experiments. All authors collaborated on the manuscript preparation.

